# Comparing sparse inertial sensor setups for sagittal-plane walking and running reconstructions

**DOI:** 10.1101/2023.05.25.542228

**Authors:** Eva Dorschky, Marlies Nitschke, Matthias Mayer, Ive Weygers, Heiko Gassner, Thomas Seel, Bjoern M. Eskofier, Anne D. Koelewijn

## Abstract

Estimating spatiotemporal, kinematic, and kinetic movement variables with little obtrusion to the user is critical for clinical and sports applications. Previously, we developed an approach to estimate these variables from measurements with seven lower-body inertial sensors, i.e., the full setup, using optimal control simulations. Here, we investigated if this approach is similarly accurate when using sparse sensor setups with less inertial sensors. To estimate the movement variables, we solved optimal control problems on sagittal plane lower-body musculoskeletal models, in which an objective was optimized that combined tracking of accelerometer and gyroscope data with minimizing muscular effort. We created simulations for 10 participants at three walking and three running speeds, using seven sensor setups with between two and seven sensors located at the feet, shank, thighs, and/or pelvis. We calculated the correlation and root mean square deviations (RMSDs) between the estimated movement variables and those from inverse analysis using optical motion capture (OMC) and force plate data. We found that correlations between IMU- and OMC-based variables were high for all sensor setups, while including all sensors did not necessarily lead to the smallest RMSDs. Setups without a pelvis sensor led to too much forward trunk lean and inaccurate spatiotemporal variables. RMSDs were highest for the setup with two foot-worn IMUs. The smallest setup that estimated joint angles as accurately as the full setup (<1 degree difference in RMSD) was the setup with IMUs at the feet and thighs. The mean correlations for joint angles, moments, and ground reaction forces were at least 0.8 for walking and 0.9 for running when either a pelvic sensor or thigh sensors were included. Therefore, we conclude that we can accurately perform a comprehensive sagittal-plane motion analysis with sparse sensor setups when sensors are placed on the feet and on either the pelvis or the thighs.

## Introduction

Gait analysis is a fundamental tool for understanding human locomotion, but is currently limited to laboratory environments. Gait analysis can include spatiotemporal, kinematic, and kinetic variables. Spatiotemporal variables provide information about the movement quality and can be used for clinical diagnoses (e.g., [1]). Kinematic, i.e., joint angles, and kinetic information, i.e., ground reaction forces, joint moments, muscle forces, as well as variables calculated from those, such as joint reaction forces, are required to understand the mechanical and physiological mechanisms of human movement. Therefore, a comprehensive analysis, including spatiotemporal, kinematic, and kinetic variables, is important in sports performance assessments (e.g., [2, 3]) and for rehabilitation (e.g., [4]). A measurement method that is cost-effective and allows for recordings “in the wild”, or outside the laboratory environment, could enable widespread use of gait analysis in different clinical and sports applications. Such methods could for example be markerless motion capture, based on video images, or motion capture using inertial measurement units (IMUs). Video-based motion capture [5, 6] still requires the person to be in the camera’s field of view. Moreover, lighting conditions, camera placement, and occlusion can affect the accuracy, limiting the flexibility of the system. IMUs are small, low cost, wearable sensors that contain an accelerometer and a gyroscope, and sometimes a magnetometer [7]. These sensors are attached to different body segments to measure their linear accelerations and angular velocities, as well as possible other signals. Since this approach is based on wearable sensors, IMU-based motion capture can be used to measure movement outside the lab in any environment [8].

While many different methods have been developed to estimate spatiotemporal variables (see [9] for a review) and kinematics (see [10] for a review) of movements from inertial sensor data, estimation of kinetics is more challenging. Joint moments are commonly computed using inverse dynamics when both kinematics and ground reaction forces (GRFs) are estimated from the IMU data [11–14]. However, this step-wise approach leads to error propagation. Errors can exist in kinematics estimates, e.g., due to soft tissue artefacts and because kinematics are usually computed by numerically integrating from linear accelerations and angular velocities to positions IMU signals, which leads to drift. Errors can also exist in GRF estimates, e.g. due to load sharing assumptions between the feet. These errors directly affect the results of the inverse dynamics calculations. Therefore, we aim to simultaneously estimate spatiotemporal, kinematic and kinetic variables from raw IMU data, which can be done using machine learning (e.g., [15–19]) or using optimal control (e.g., [2, 20]). Machine learning models can be trained to directly map IMU data to biomechanical variables (e.g., [15–19]). However, an important limitation of machine learning models is that it cannot be proven that the resulting machine learning models outputs dynamically consistent results, meaning that the kinematic and kinetic variables might not follow the laws of physics. Furthermore, machine learning models are typically trained and tested using lab-based optical motion capture (OMC) and IMU data, which means that there is no proof that such a model will perform well “in the wild”. Another major challenge when applying machine learning to estimate biomechanical variables is the availability of training data, since the machine learning model accuracy has been shown to improve with the size of the training dataset [21].

In order to obtain interpretable and dynamically consistent motions, we have developed an optimal control approach to simultaneously estimate the kinetics and kinematics of walking and running based on raw accelerometer and gyroscope measurements [22]. With this approach, we find a dynamics simulation for a musculoskeletal model, such that virtual inertial sensor data of the simulation match the recorded inertial sensor data as closely as possible. This approach does not require a training dataset, inherently overcomes the aforementioned drift problems, and can mitigate soft-tissue artifacts [22]. An optimal control problem is solved to find a simulation for a sagittal plane musculoskeletal model that minimizes effort while also minimizing a tracking error between virtual and measured inertial sensor data, specifically angular velocities and linear accelerations. The resulting simulation contains spatiotemporal, kinetic, and kinematic variables that are dynamically consistent [23]. Since we use a musculoskeletal model, the kinetic variables are the ground reaction forces, joint moments and muscle forces. We include muscle dynamics to constrain torques to be physiologically realistic. Furthermore, the tracking approach avoids integration of the inertial sensor measurements and thus the related integration drift. We previously showed that with this approach, we could estimate joint angles and joint moments similar to those obtained through inverse kinematics and inverse dynamics based on OMC and GRF measurements [22].

However, estimating both kinetics and kinematics from inertial sensor data commonly requires a sensor on each body segment of interest. For example, to estimate leg kinematics and kinetics using optimal control, we used 7 inertial sensors [22]. Others used 8 sensors to include the trunk [24], or 17 sensors for a full-body analysis [12]. When more sensors are placed on the body, the wearer, and thus their motion, can be increasingly affected [25]. Furthermore, using too many sensors increases cost, reduces practicality, and increases the chance of errors in IMU placement [26]. Therefore, our aim is to create an IMU-based motion analysis approach that is as unobtrusive as possible by reducing the number of necessary sensors. Sparse sensor sets have been developed to estimate individual gait variables, such as spatiotemporal (e.g., [1]), kinematic [26–29], or kinetic [30] variables. Furthermore, neural networks have been investigated to estimate specific quantities of interest using only two IMUs on the shanks [26] or the feet [1, 28]. Others have proposed physics-based optimization to track the sensor orientation with a torque-driven model [30] or a human body shape model [29]. However, these approaches do not provide a comprehensive analysis including various biomechanical variables, or neglect physical correctness or muscle dynamics. Furthermore, to our knowledge, no systematic comparison has been performed to evaluate different sensor configurations for estimating spatiotemporal variables, kinematics, and kinetics during walking and running.

In this work, we performed such a systematic comparison to evaluate how well spatiotemporal variables, joint moments, ground reaction forces, and joint angles can be estimated from sparse sensor setups using an optimal control approach. Previously, we developed an optimal control approach for a full lower-body sensor setup [22]. Since realistic walking simulations of musculoskeletal models can also be created by solving optimal control problems with no data tracking [31], we expect that we can also apply the approach developed by [22] to sparse sensor setups as well. Here, we therefore investigated the accuracy of reconstructive simulations created by solving optimal control problems for six sparse sensor setups and the full setup. We created the six sparse sensor setups by varying the number of IMUs included, as well as the body segments on which they were attached. We used a minimal setup of two IMUs placed on the feet, three setups with sensors on two locations: feet and shanks, feet and thighs, and feet and pelvis, and two setups with sensors at three locations: feet, shanks and pelvis and feet, thighs and pelvis, and compared those to a full setup with sensors on the feet, shanks, thighs and pelvis. We then created optimal control simulations of walking and running for each sensor setup and investigated the difference between the resulting spatiotemporal variables, i.e., speed, stance time, and stride length, gait kinematics, i.e., joint angles, and gait kinetics, i.e., joint moments and ground reaction forces and those obtained using OMC and using the full lower-body sensor setup as used by [22].

## Methods

We reconstructed walking and running motions of a sagittal-plane musculoskeletal model using raw inertial sensor measurements, i.e., raw gyroscope and accelerometer data, from six different sparse sensor setups and one full lower-body sensor setup (see Fig. 1). Sensors were placed symmetrically for all setups, and each setup included sensors at the feet. We included feet sensors in all setups, since these sensors can be attached to the shoe, and are therefore unobtrusive, while they have also provided reliable information in past studies [1]. For each sensor setup, we created musculoskeletal model simulations by solving optimal control problems that minimized the difference between measured and virtual sensor data. To evaluate the different sensor setups and their respective simulations, we compared the difference between these simulations and an OMC analysis.

**Fig 1.**
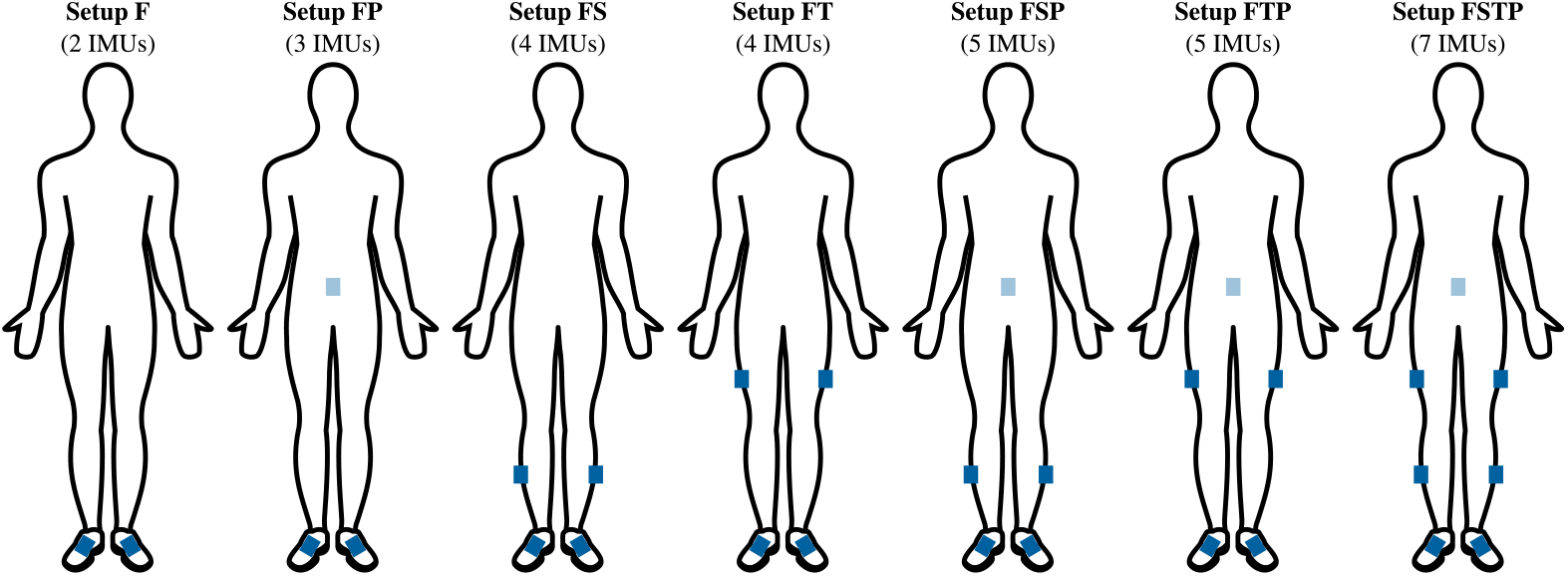
Seven sensor setups. The blue rectangles indicate the approximate location of the inertial measurement units (IMUs). The sensor setups are abbreviated using the first letter of the segments equipped with a sensor: F-feet, S-shanks, T-thighs, P-pelvis. The setup FSTP corresponds to a full lower-body sensor setup with seven IMUs.

## Experimental Data

We used measured data previously recorded with seven custom-built IMUs (Portabiles HealthCare Technologies GmbH, Erlangen, Germany) placed on the pelvis, legs and feet (Fig. 1, setup FSTP) and an OMC system including one force plate. In [22], we described the data recording and data pre-processing and used this dataset to evaluate optimal control simulations from the full lower-body sensor setup (setup FSTP). In this work, we evaluated six sensor setups with sparser sensor placement (Fig. 1, setup F, FP, FS, FT, FSP, and FTP), and compared those to the full lower-body sensor setup as using in [22] and OMC data. The raw and pre-processed data (mean and standard deviation over 10 trials) are available in [32].

This dataset contained recordings of 10 healthy male participants (age: 27.1 *±* 2.6 years, height: 181.9 *±* 5.3 cm, weight: 76.9 *±* 8.6 kg) at six different speeds (slow walking: 0.9-1.0 m s^−1^; normal walking: 1.3-1.4 m s^−1^; fast walking: 1.7-1.8 m s^−1^; slow running: 3.1-3.3 m s^−1^; normal running: 3.9-4.1 m s^−1^; fast running: 4.7-4.9 m s^−1^).

Participant recruitment and data collection took place from June to August 2016. All participants gave their informed written consent prior to participation. The study was conducted in line with the ethical principles of the Declaration of Helsinki and it was approved by the ethics committee of the Medical Faculty at the FAU Erlangen-Nürnberg, Germany (Ref.-No.: 106 13 B).

As described in [22], IMU axes were aligned with the body segment axes using functional calibration movements. From the OMC-data, sagittal plane joint angles and joint moments were calculated using the GaitAnalysisToolkit (see [33]) according to [34]. IMU and OMC measurements were synchronized using a custom flash trigger system. In addition, we corrected for a small time offset (approx. 20 ms) between IMU and OMC measurements, which was determined by calculating the cross-correlation between the time derivative of the hip, knee, and ankle joint angles obtained from the OMC analysis and the angular velocity measured by the adjacent gyroscopes.

## Musculoskeletal Model and Dynamics

We used a sagittal-plane musculoskeletal model to create our gait simulations [22, 35]. The skeleton consisted of seven segments: head-arms-trunk (HAT), two upper legs, two lower legs, and two feet, and had nine kinematic degrees of freedom **q**: the position and orientation of the trunk, two hip angles, two knee angles, and two ankle angles (Fig. 2). We personalized the musculoskeletal model’s segment masses, lengths, center of mass locations and moments of inertia based on the study participants’ full-body height and weight [34]. We modeled 16 lower-leg muscles as three-element Hill-type muscles [36] (Fig. 2). Overall the model’s state, **x**(*t*), was described by the nine degrees of freedom **q**(*t*), the corresponding nine velocities **v**(*t*), 16 contractile element lengths **l**_**CE**_(**t**), and 16 muscle activations **a**(*t*): **x**(*t*) := [**q**(*t*) **v**(*t*) **l**_**CE**_(*t*) **a**(*t*)]^*T*^ for 0 ≤ *t* ≤ *T* with movement duration *T* .

**Fig 2.**
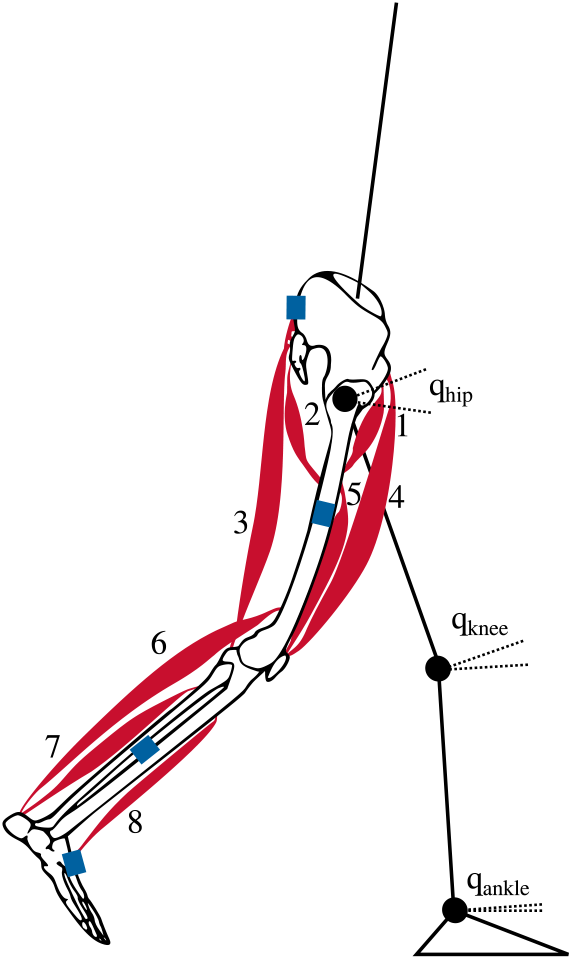
Musculoskeletal model with seven rigid segments and 16 Hill-type-muscles, eight per leg: 1 – iliopsoas, 2 – glutei, 3 – hamstrings, 4 – rectus femoris, 5 – vasti, 6 –gastrocnemius, 7 – soleus, and 8 – tibialis anterior [22, 35]. The blue rectangles indicate the approximate location of the inertial measurement units (IMUs).

We modelled contact between the musculoskeletal model and the ground using two contact points at each foot (heel and toe) [22]. We used a penetration-based contact model to describe the vertical ground reaction force, and a friction model for the horizontal ground reaction force. The contact model equations are further described in [37]. Each contact point was described by its global (*x, y*) position, and the anterior-posterior (*F*_*x*_) and vertical (*F*_*y*_) force.

We defined the dynamics by combining the musculoskeletal model and the contact model. The model was controlled through the 16 muscle excitations, **u**(*t*) for 0 ≤ *t* ≤ *T* [22]. The system dynamics were fully described by implicit differential equations as a function of 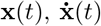 and 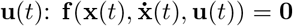. The system dynamics, including multibody dynamics, muscle-tendon equilibrium, activation dynamics, and contact model, **f** (), were twice differentiable with respect to all 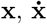 and **u** [36], such that the optimal control problems can be solved using a gradient-based optimization algorithm.

We calculated virtual IMU data using a virtual sensor model that was added to the musculoskeletal model [22]. We placed virtual sensors in their respective position, as measured during the experiment, on the musculoskeletal model, and calculated the virtual sagittal plane accelerometer and gyroscope data from the model state and its derivative [22]. The gyroscope signal represented the angular velocity of the respective body segment relative to the global coordinate system and was derived from the generalized velocities. The acceleration signal represented the linear acceleration at the respective sensor position on the body segment relative to the global coordinate system and was a combination of the acceleration of the body segment origin and the acceleration caused by the segment rotation. The used equations can be found in [22].

### Optimal Control Problems

We created gait simulations from inertial sensor data by solving optimal control problems. In these optimal control problems, a multi-objective optimization was solved to find a periodic walking or running cycle that minimized muscular effort, while also tracking the measured inertial sensor data. We constructed the tracking objective to minimize the squared difference between measured accelerations and angular velocities from the IMUs and the corresponding simulated accelerations and angular velocities from the virtual sensor model, averaged over the duration of a gait cycle *T* :

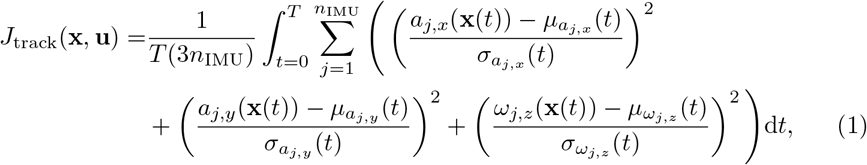

where the number of IMUs *n*_IMU_ ranged between two and seven depending on the setup in Fig. 1. Since we used a two-dimensional musculoskeletal model defined in the *x*-*y*-plane, we calculated the virtual accelerations in *x*- and *y*-direction *a*_*j*,*x*_ and *a*_*j*,*y*_, and the angular velocity *ω*_*j*,*z*_ around the *z*-axis for each IMU *j* ≤ {1, …, *n*_IMU_} [22]. We tracked experimental IMU data *µ* averaged over ten gait cycles. The squared difference between simulated and measured signals was divided by the variance of the measured data *σ*^2^. This approach leads to a smaller weight being applied when there is a large variance, e.g., due to soft tissue artefacts, and automatically provides an appropriate weighting between different sensor modalities.

We computed muscular effort as squared muscle excitations averaged over the duration of a gait cycle *T* and normalized to the squared walking or running speed *v*

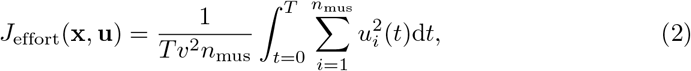

where *n*_mus_ = 16 corresponds to the number of muscles. For a faster convergence, we added a regularization term that minimized the squared rate of change of all states and all muscle excitations [38, 39]. These objective terms yielded the following optimal control problem [22, 36]:

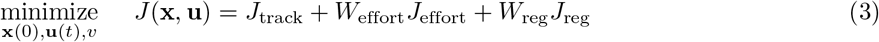

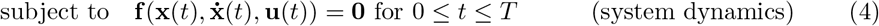

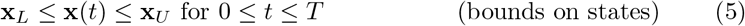

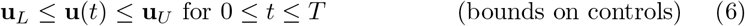

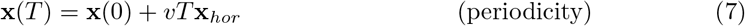

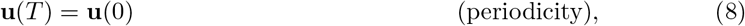

where *W*_*eff*_ = 300 and *W*_reg_ = 10^−5^ are the weightings of the effort term and regularization term, respectively. In Eq. 7, we enforced a periodic forward motion, where *x*_*hor*_ is a vector of states to which a horizontal displacement applies, which are the horizontal pelvis position and the horizontal contact point positions.

To solve the optimal control problems, we transcribed them (objectives and constraints) using direct collocation with a backward Euler discretization and *N* = 100 time nodes [22, 36]. We added an extra collocation node *N* + 1 to evaluate our periodicity constraints (Eq. 7 and Eq. 8), such that the state and input at node *N* + 1 should be the same as at node 1 with a horizontal translation. Our decision variables consist of the model’s state, **x**, and input, **u** at each time node, as well as the global contact point locations and forces. We chose to include these, since preliminary work showed that this approach speeds up the optimization. We then also added constraints to ensure that the location and forces match those calculated from the contact model using the model’s state. Furthermore, our decision variables also included the motions’s speed and duration. In total, 8284 variables (101 time nodes *·* (18 multibody states + 32 muscle states + 16 muscle stimulations + 4*·*4 contact point decision variables) + speed + duration = 8284) were optimized with 6682 constraints (100 time nodes *·* (18 multibody dynamics equations + 32 muscle dynamics equations + 4*·*4 contact model equations) + 82 periodicity constraints = 6682). We used the same initial guess and the same objective weights as [22]. We solved the resulting large scale nonlinear optimization problems with IPOPT with MUMPS [40]. We solved 420 optimal control problems, i.e., simulations for seven setups for 10 participants at six speeds, on a computer cluster using one Intel Xeon E3-1240 v6 for each simulation.

### Data Analysis

We analyzed the convergence and computation time of the optimal control problems, followed by a comparison of the spatiotemporal variables (walking speed, stance time, stride length), the kinematic variables (sagittal plane joint angles), and the kinetic variables (sagittal plane joint moments and ground reaction forces) as calculated with the OMC measurements and the IMU measurements. First, we determined the speed as the translation of the right heel divided by the duration of the gait cycle, the stance time as the ratio of the duration of the stance phase [41] and the duration of the gait cycle, and the stride length as the translation of the right heel. We quantified the similarity of the spatiotemporal variables between the different IMU setups and the OMC setup using the root mean square deviation (RMSD). For each walking and running speed, we calculated the RMSD between the results calculated with the different IMU setups and the results of the OMC analysis using the data of all participants. In addition, we generated scatter plots comparing the spatiotemporal variables obtained from the OMC measurements with those obtained from the different IMU setups in order to observe systematic differences between the setups. Next, we compared kinetic and kinematic trajectories of the right leg between the results calculated with the different inertial sensor setups and the results of the OMC analyis using the coefficient of multiple correlation (CMC) [42] and RMSD. We calculated the CMC and the RMSD for each walking or running cycle of each participant individually and then determined the mean (using Fisher’s Z-transform for the CMC) over all walking and running cycles and participants. We computed the CMC and RMSD for the entire gait cycle for the joint angles and for the stance phase for the joint moments and GRFs.

## Results

We solved 420 optimizations, which required a mean *±* standard deviation CPU time of 48 *±* 3 min (Table 1, Supplementary file S01). 418 of the 420 simulations converged, while no optimal solution was found for two simulations. For one participant, the restoration phase failed for setup FP in the slow walking trial, while for another participant, this happened for setup F in the fast walking trial. We removed these two simulations from the analysis. CPU times for walking simulations were higher than for running simulations, but similar for the different setups except for setup F for walking. We added the measured IMU trajectories and those calculated from the optimal control simulations in supplementary files. S01 includes averaged results for walking and running, and S02 includes individual results for each trial, participant, and sensor setup.

**Table 1.** CPU time for converged optimizations of the different inertial sensor setups (mean ± standard deviation).

| Setup | F | FP | FS | FT | FSP | FTP | FSTP |
| --- | --- | --- | --- | --- | --- | --- | --- |
| CPU time in min for walking | $110 \pm 85$ | $77 \pm 33$ | $61 \pm 28$ | $54 \pm 20$ | $63 \pm 28$ | $63 \pm 22$ | $68 \pm 30$ |
| CPU time in min for running | $27 \pm 10$ | $26 \pm 9$ | $27 \pm 9$ | $31 \pm 11$ | $28 \pm 9$ | $29 \pm 10$ | $34 \pm 10$ |

We found that the RMSDs of the spatiotemporal variables were similar between all sensor setups, though setup F performed worst (see Fig. 3). The RMSDs for speed and stride length were highest for setup F (speed^1^: [0.15; 0.55] m*/*s; stride length:

**Fig 3.**
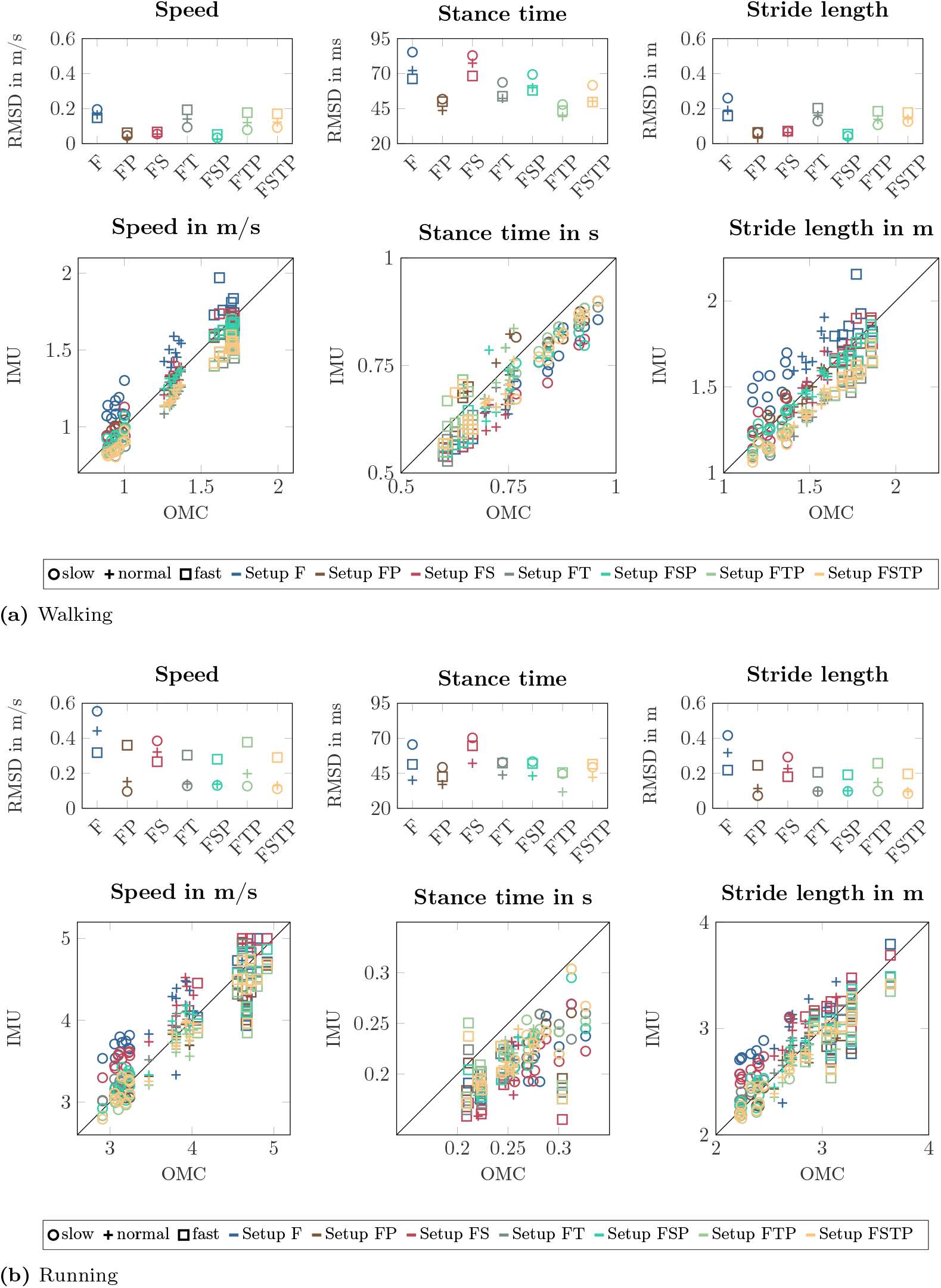
Root mean square deviation (RMSD) and scatter plots for speed, stance time, and stride length between the different inertial measurement unit (IMU) setups and the optical motion capture (OMC) for (a) walking and (b) running. We calculated the RMSDs over all participants for each walking and running speed and each setup. The scatter plots show the spatiotemporal variables for each walking and running cycle of each participant, where the thin black lines represent a perfect match between IMU and OMC data.

[0.16; 0.42] m) for most walking and running conditions. When adding a pelvis sensor (setup FP), RMSDs of speed and stride length decreased (speed: [0.03; 0.36] m*/*s; stride length: [0.04; 0.24] m). The setups without a sensor on the thighs (setups FP, FS, and FSP) resulted in lower RMSDs for speed and stride length. Speed and stride length were systematically underestimated when a thigh sensor was added (see Fig. 3). For the setups with sensors only on the lower leg segments (setups F and FS), the stance time RMSD was worst, up to 27.2 ms higher than the full sensor setup (setup FSTP).

When visualizing the mean of our simulations for each setup with stick figures, we found that the simulations without a pelvis sensor had more forward trunk lean than the full sensor setup (Fig. 4). We also observed that the step length for setup F is larger than the step length of all other setups for walking and running, supporting the larger RMSDs shown in Fig. 3 for the stride length. This bias is also evident in the stride length correlation plots for setup F in Fig. 3. Furthermore, for walking, a difference in knee flexion angle is visible in the trailing leg, where the solutions with thigh sensors have a larger knee flexion angle than those without.

**Fig 4.**
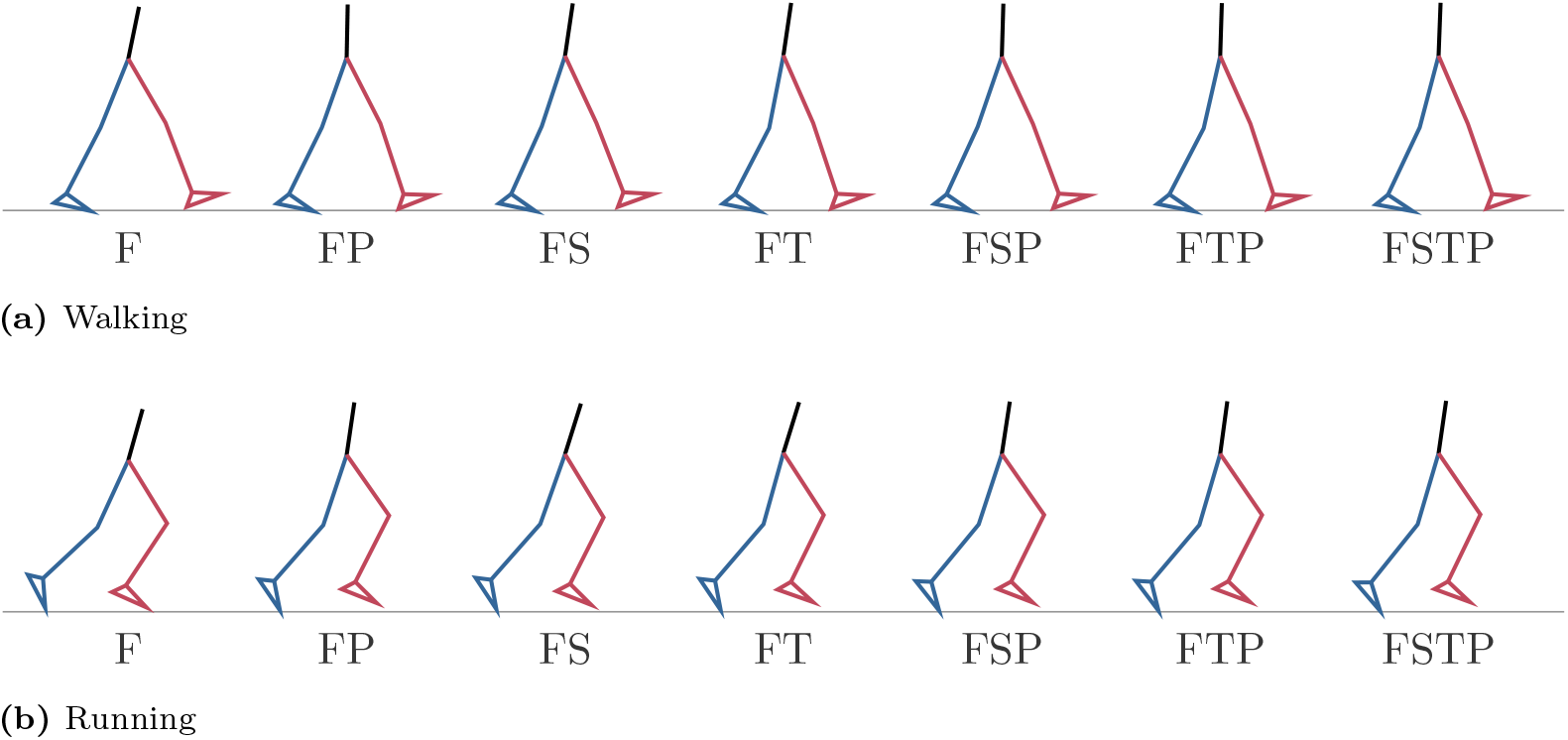
Stick figure representing the mean kinematics at the first time point of the different inertial measurement unit (IMU) setups for (a) walking and (b) running over all participants. The stick figures show that without a pelvis sensor (setups F, FS, and FT), forward trunk lean is too large.

Our simulations showed that joint angle, joint moment, and ground reaction force estimates generally benefitted from having sensors placed at the pelvis or thighs in addition to those at the feet (see Fig. 5, supplementary file S01 for individual results). For setup F, knee flexion was too high during stance for both walking and running, and so was the peak knee extension moment. For running, the peak ankle dorsiflexion angle was too high as well. For setups without thigh sensors (F, FP, FS, and FSP), we also observed that the joint range of motion of the hip was larger than in the OMC result for the walking simulations, and so was the knee extension during late stance for the walking simulations. A similar larger peak flexion moment was observed in the knee. The forward progression of the ankle dorsiflexion was too fast for all setups except setup FS for the walking simulations, while a similar fast progression was observed in the ankle plantarflexion moment for setups F, FP, FT, and FS, both for walking and running. The peak hip flexion moment occurred later in the gait cycle for setup FS for both walking and running, and for setup FT for running. The peak hip flexion angle in the running simulations was larger than for the OMC result in the setups F, FT, and FS, while the peak extension moment is larger than in the OMC results for all setups except FT. Furthermore, the anterior-posterior GRF displayed larger braking and push-off forces for all IMU setups, most prominently for setup F, while the peak vertical GRF was too high for setups F and FS.

**Fig 5.**
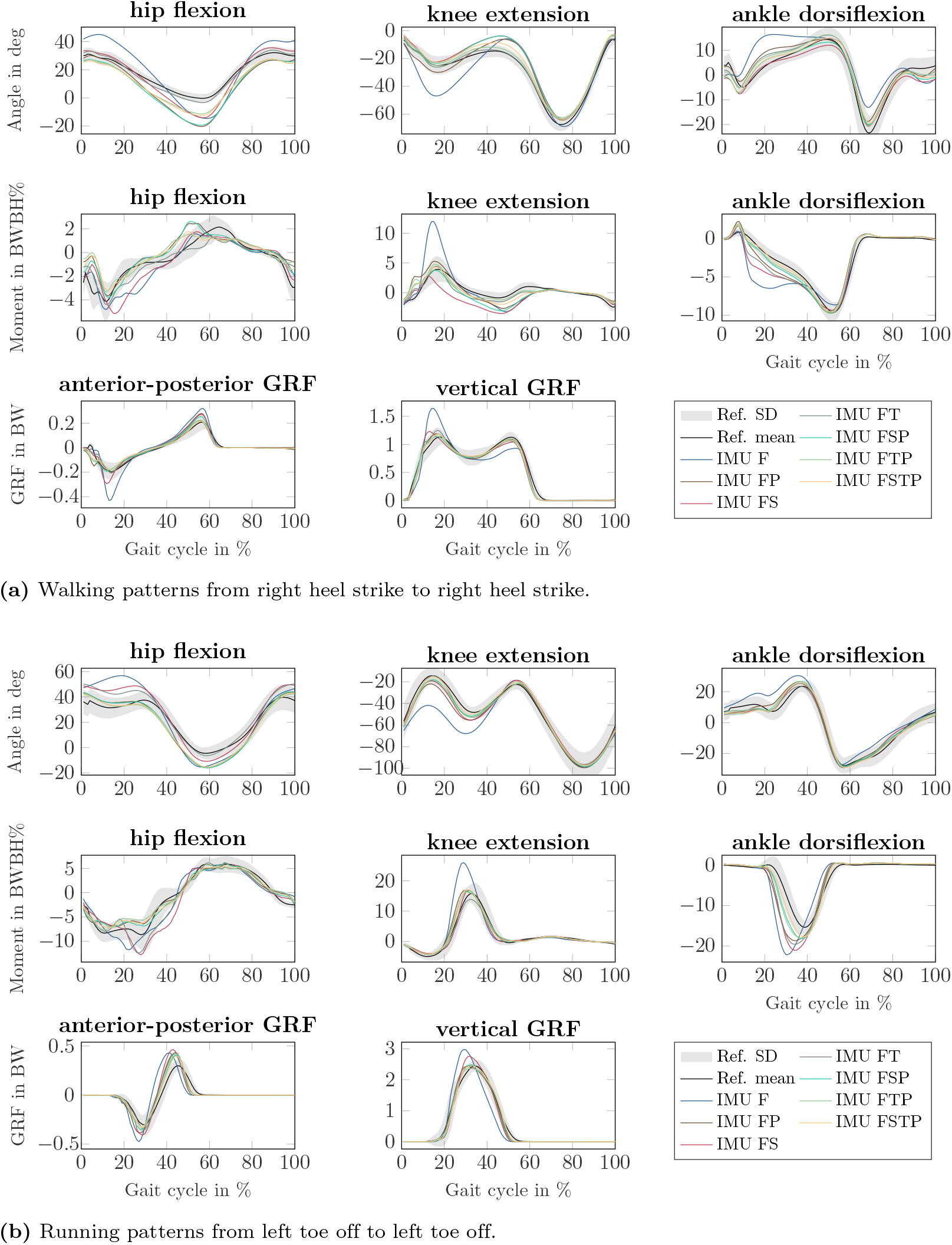
Sagittal-plane joint angles, joint moments, and ground reaction forces (GRFs) of the right lower limb for (a) walking and (b) running at all speeds, from the different inertial measurement unit (IMU) setups (colored lines) and the references values from optical motion capture system and force plate data (mean: black line, standard deviation: grey fill). All lines represent the mean over all participants. Joint moments were scaled to bodyweight bodyheight percent (BW BH %) and GRFs were scaled to bodyweight (BW).

We found that the change in CMCs with the sensor setup varied between the different kinematic and kinetic variables, and that the difference in CMC was generally larger between setup FSTP and setup F than between setup FSTP and all other sparse sensor setups (Fig. 6). The mean CMCs of the joint angles, ankle joint moment, knee joint moment for running, and GRFs was lowest for setup F, while the mean CMC of the knee moment for walking and of the hip moment for running was lowest for setup FS. The mean CMC of the hip moment for walking was even lower than that of setup F for setups FSTP, FTP, FSP, and FP, and higher only for setups FS and FT. The largest difference in mean CMC between setup F than setup FSTP was 0.14 (running, anterior-posterior GRF). The largest difference in mean CMC between setup FSTP and each of the other sparse setups was at least 50% smaller, up to 0.07. Only for setup FS, the largest difference to setup FSTP (walking, knee moment) was 100% larger than the largest difference between setup F and setup FSTP (0.29 vs. 0.14). For some variables and sparse setups, the mean CMC was larger than for setup FSTP, specifically the mean CMC of the hip and ankle angle and hip moment for walking in setup FS, the mean CMC of all joint angles, the hip moment, and the anterior-posterior GRF for walking and of the hip and ankle angle for running in setup FT, the mean CMC of the ankle angle and moment for walking in setup FSP, and the mean CMC of all joint angles and the ankle moment for walking and of the vertical GRF for walking and running in setup FTP. Furthermore, while the range of CMC values was small for the joint angles, ankle moments, and GRFs, there was a larger variation for the hip and knee joint moment, where some simulations did not correlate with the OMC result at all.

**Fig 6.**
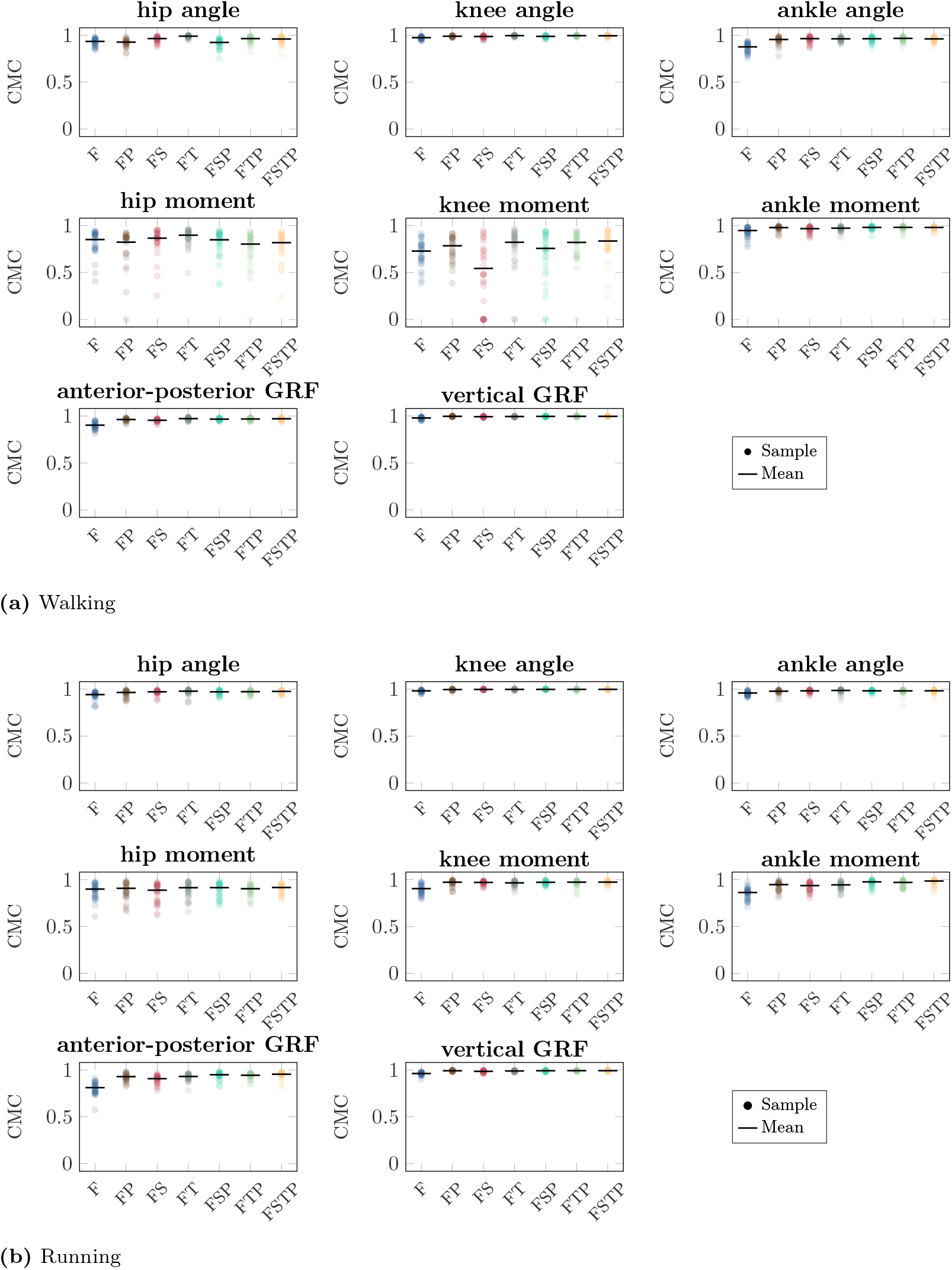
Coefficient of multiple correlation (CMC) for the sagittal-plane joint angles, joint moments, and ground reaction forces (GRFs) between the different inertial measurement unit (IMU) setups and references values from optical motion capture system and force plate data for (a) walking and (b) running. The circles show the CMC for each walking and running cycle of each participant. The bar shows the mean value over all cycles and participants computed using Fisher’s Z-transform.

Our RMSDs showed the lower accuracy of setup F more clearly than the CMC, while the setups with sensors on the thighs or on the pelvis were most accurate for all variables (Fig 7). Similar to the CMC, the RMSD of all variables was highest for setup F, except for the hip moment for running, where setup FS yielded the highest RMSD. The joint angle RMSD was up to 11 deg higher (running, knee) for setup F than setup FSTP, while the GRF RMSD was up to 0.37 BW higher (running, vertical) and the joint moment RMSD up to 5.5 BW BH % higher (running, ankle). The differences between setup FSTP and the other sparse setups were smaller than those between setup FSTP and setup F. Compared to setup FSTP, setups FP and FSP had the largest increase in joint angle RMSD with 3 deg (hip, walking), while the largest increase was 2.6 deg (walking, knee) for setup FS and 0.8 deg for the setups FT and FTP (running, knee). The joint angle RMSD was even lower than for setup FSTP for setups FS (walking, ankle), FSP (walking and running, ankle), FT (walking, hip and knee, and running, ankle), and FTP (walking, all joints). The GRF RMSD for setup FSTP was up to 0.035 BW lower (running, anterior-posterior) than for setup FP, up to 0.051 BW lower (running, vertical) than for setup FT, up to 0.13 BW lower (running, vertical) than for setup FS, and up to 0.015 BW lower (running, vertical) than setups FSP and FTP. The GRF RMSD was even lower for setups FP (walking, vertical), FT (walking, anterior-posterior), and FTP (running, vertical) than for setup FSTP. The joint moment RMSD was consistently highest in the ankle for running. Compared to setup FSTP, it was up to 2.6 BW BH % higher for setups FP and FT, up to 3 BW BH % higher for setup FS, up to 1.2 BW BH % higher for setup FTP, and up to 0.8 BW BH % higher for setup FSP. The joint moment RMSD for setup FT (walking, hip and knee) was even lower than for setup FSTP. The range of the hip and knee moment RMSD of were similar to those of the ankle moment, while for the CMCs, the difference in range between the hip and knee moment and the ankle moment was larger.

**Fig 7.**
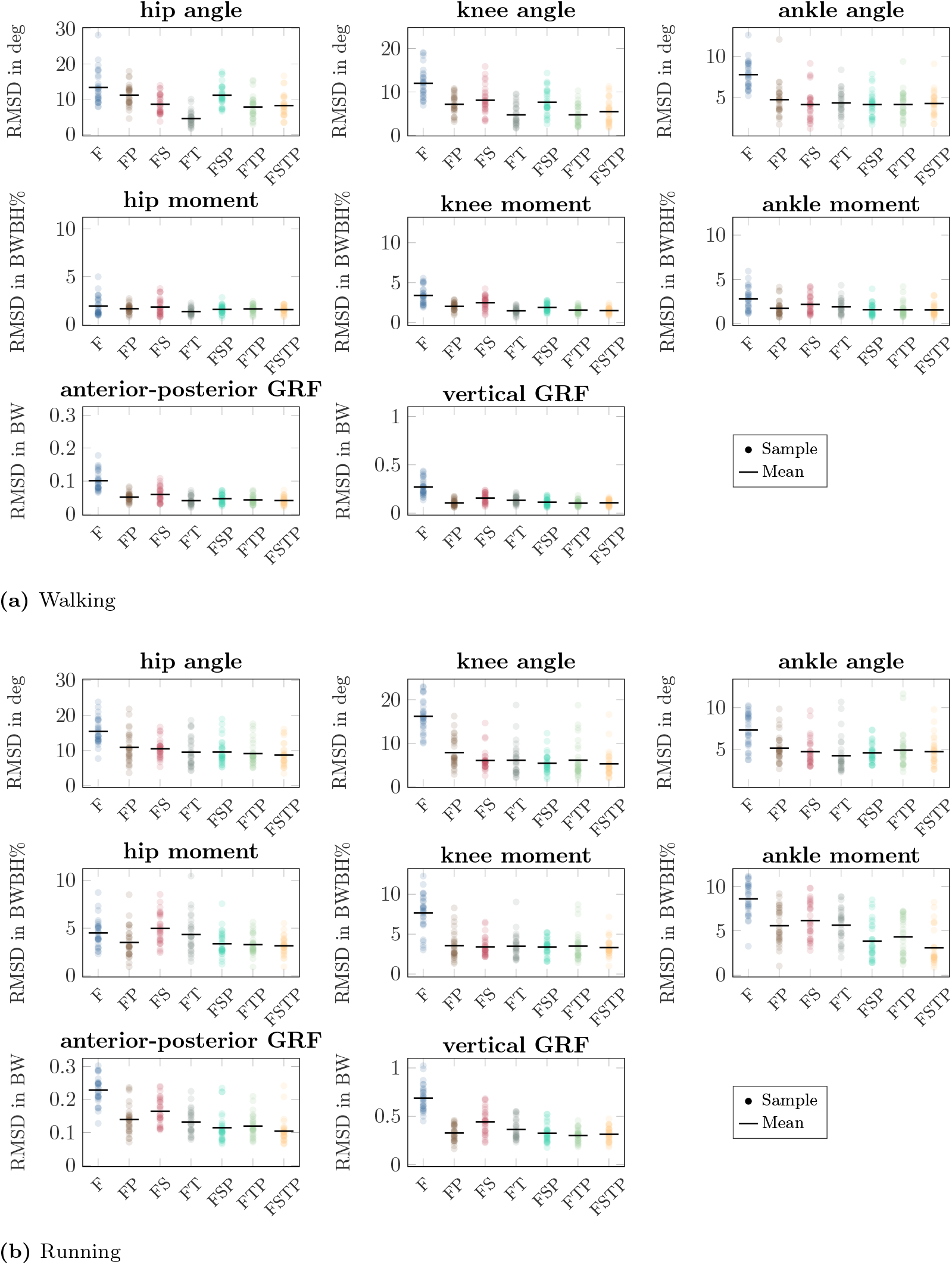
Root mean square deviation (RMSD) for the sagittal-plane joint angles, joint moments, and ground reaction forces (GRFs) between the different inertial measurement unit (IMU) setups and references values from optical motion capture system and force plate data for walking and (b) running. The circles show the RMSD for each walking and running cycle of each participant. The bar shows the mean value over all cycles and participants. Joint moments were scaled to bodyweight bodyheight percent (BW BH %) and GRFs were scaled to bodyweight (BW).

## Discussion

We evaluated optimal control simulations from six different sparse IMU setups by comparing their accuracy against a full sensor setup and a OMC-based motion analysis. We showed that it is possible to reconstruct sagittal-plane walking and running biomechanics, including spatiotemporal, kinematic and kinetic variables, from sparse inertial sensor setups. We found that the correlations between the IMU-based analysis and the OMC-based analysis were large for all sensor setups with mean CMC values ranging between 0.81 and 1 for joint angles, ankle moment and GRF, while mean CMCs of the hip and knee moment ranged between 0.54 and 0.97. Furthermore, we found that the sensor setups that included sensors on either the thighs or the pelvis only had small performance drops compared to FSTP, while the differences were much larger for setup F for all tested spatiotemporal, kinematic, and kinetic variables and for setup FS in stance time and kinetics, though the results depended on the variable type.

When comparing different sparse IMU setups, we found that in general it is beneficial to use more than only two sensors at the feet, since RMSD improved or remained similar for the spatiotemporal variables, and so did the RMSD and CMC for most kinetic and kinematic variables (Fig. 3, Fig. 7, and Fig. 6). We also found that the optimal sensor setup depends on the application variables of interest and that a full sensor setup does not always produce the best results. For the spatiotemporal variables, especially for walking, adding sensors to the thighs reduced prediction accuracy (Fig. 3), while adding thigh sensors improved similarity of the kinetic and kinematic patterns (Fig. 5). Adding a sensor to the pelvis also improved the accuracy of kinematics and kinetics, but not as much as when using thigh sensors. Omitting shank sensors seems reasonable when sensors at the feet and the pelvis or thighs are used since estimation accuracy either only slightly improved or even decreased when adding shank sensors.

We compared different sparse sensor setups to previous work and found higher or similar RMSDs in our work. Previously, we compared the results of the full setup [2], and showed that kinematics and kinetics were comparable to Karatsidis et al. [12].

Weygers et al. [10] reviewed kinematics estimates from inertial sensors of gait and other movements. We found that our kinematic estimates for all setups are similar to other gait estimates (e.g., [43]). Similar to our work, RMSD was generally highest for the hip joint angle than and lowest for the ankle joint angle [10, 43]. Furthermore, the mean RMSD for the knee angle and vertical GRF of setup FP (11 *±* 4 deg and 0.3 *±* 0.1 BW, respectively) are comparable to those of a similar setup reported in [44] (5 − 20 deg for the peak knee extension angle and 0.2 − 1.3 BW for the peak vertical GRF, who trained several neural networks to output specific kinematic and kinetic variables. These comparisons highlight that our approach using optimal control is as accurate as other commonly used approaches, while it allows for different biomechanical variables to be estimated simultaneously. However, inference of machine learning models is generally fast, while our simulations require half an hour to two hours to solve.

The estimated joint moments of the reconstruction were surprising, specifically the large range in CMC for the hip and knee moment. The large range in CMC for the hip and knee moment can be explained by the differences between the processing of the OMC and IMU data. The processing of the OMC data included filtering, followed by inverse kinematics and inverse dynamics [34] with a different underlying model. It might be that the low-pass cut-off frequency in our filter was too high, allowing for more high frequency signal to pass than feasible in the optimal control simulation. This would especially affect the CMC but not the RMSD, as we observed in our results.

Furthermore, the general differences in processing yield that the kinetics and kinematics of the OMC result cannot be reproduced exactly in an optimal control simulation. This comparison is therefore challenging, since the exact ground truth is unknown, as joint moments and joint angles cannot be measured directly. To remove most differences between the IMU- and OMC-based analysis, and make the comparison as direct as possible, the optimal control simulations tracking IMU data could be compared against optimal control simulations tracking marker and GRF [23].

Furthermore, the setup of the optimal control problem used for the biomechanical gait reconstructions could also affect the accuracy of the simulations. For example, contact between the ground and our musculoskeletal model is created by a rigid foot with two penetration-based contact points, one at the heel and one at the toe, which is a simplification of foot-ground contact as it happens in practice. These model simplifications cause different kinetics to be associated with certain kinematics for the musculoskeletal model than for the experimental participants, such that the kinetics as recorded by and estimated from OMC cannot be achieved. The effect of this foot-ground model on the ankle moment accuracy was already observed for the full lower-body sensor setup [22].

Another choice in the optimal control setup is the weighting between the effort and tracking objective. We chose our weighting based on [22] and normalized it to the number of sensors to ensure that the objective value was similar for all setups. This weight term could still be tuned individually for each sensor setup, and could even be tuned for each sensor. For example, a larger weight on the pelvis or thigh sensors could improve accuracy, especially in a sparse sensor setup, because we found that its inclusion in the sensor setup is important. Additional objective terms could be considered in future work to compensate for missing information in the IMU data. For example, Falisse et al. found that a multi-objective cost function minimizing metabolic energy rate, muscle activations, and joint accelerations predicted human-like walking patterns without tracking measured data [31].

There are also different approaches to track data in the optimal control problem. Here, we chose to track the mean and variance over 10 gait cycles for each participant. Instead, we could have also created simulations for each gait cycle individually [20].

This decision depends on the motivation for creating the simulations. When the goal is to analyze a longitudinal parameter, such as repeated loading or fatigue, the benefit of tracking an average gait cycle is that the simulation does not reflect natural variation. From a technical perspective, averaging and then simulating is also faster than first simulating and then averaging. An additional advantage is that we adjust the weighting to the natural variance of the movement, since a large variance reduces the objective weighting and a small variance increases it. Due to this, parts of the movement with little variance are tracked more closely that parts with much variance. However, when the goal is to provide biofeedback, for example during training, it makes more sense to provide this for each gait cycle individually and therefore simulate individual gait cycles.

A further choice in the optimal control problem is the transcription method. Here, we chose a first-order backward Euler discretization to transcribe the objective and constraints of the optimal control problem. Backward Euler is known to add artificial damping to the solution, meaning that the motion requires more energy. Previous work [2, 36] showed that this error is systematic, which allows for comparisons between two simulations created with this approach. A further uncertainty might appear from the discretization error. We performed a convergence analysis and did not find a difference between the simulations with 100 time nodes and the same simulations with 200 and 400 time nodes. Still, a higher-order method [45, 46] would decrease the discretization error for the same number of time nodes. It has not yet been investigated how the (order of) the transcription method affects the accuracy of a simulation with respect to experimental data. We expect that this error is smaller than the error that can be expected due to uncertainty in the musculoskeletal model parameters, where different sources can lead to a difference of up to 20% in joint moments [47].

We have shown that it is possible to create sagittal-plane reconstructions of walking and running, addedand estimate kinetics and kinematics from sparse sensor setups We were able to create reconstructions with only three sensors (on the feet and the pelvis) or only four sensors (on the feet and the thighs) that had only a few noticeable differences to those created with a full lower-body sensor setup. These results imply that the improved usability of having only three or four sensors does not lead to a considerable performance drop of the movement reconstructions. Our results support previous work that showed that, while such movement reconstructions are possible using only foot sensors, results are worse in this approach [22]. In contrast, deep neural networks that directly map foot-worn sensor data to lower-leg kinematics have shown promising results [28]. They reported mean RMSDs for sagittal-plane joint angles of 4 to 5 deg, whereas our tracking simulations using only foot-worn IMUs resulted in mean RMSDs of 4 to 13 deg. One interesting direction for future work would be to combine deep learning and physics-based optimization to obtain physically correct human motion from deep neural networks [19, 48].

Different sensor setups could be investigated to further optimize both usability and accuracy. For example, a sensor setup with three sensors on the shanks and the pelvis could be considered. We chose the setup with sensors on the feet to include as many body segments as possible between the sensors, because no information would be available about the feet when sensors are placed on the shank. Furthermore, it might be possible to increase sparsity by placing sensors asymmetrically on different body segments when symmetric motions are recorded, to avoid double recording of the same signal, e.g., on the left shank and right thigh. Such setups could be further investigated using an observability analysis to investigate the exact working conditions in which kinematic and kinetic variables can be inferred from simulated inertial measurement data. So far, such an analysis has been performed using recurrent neural networks (RNNs) to estimate the observability of kinematics on simplified mechanical linkages [49], as well as for sparse inertial motion tracking [50]. Nevertheless, in future work such an RNN-based observability analysis might be used to study the observability of both kinematic and kinetic variables in sparse IMU setups for realistic biomechanical kinematic chains, such as we used here.

These results will help those interested in studying motion “in the wild”, or outside of the lab environment, since we have shown that using sparse sensor setups will not necessarily reduce accuracy of the resulting simulations. Sparse sensor setups with high accuracy enable capturing of real-life data with reduced burden on patients compared to a full-sensor setup. This real-life data can then support decision-making by providing objective data complementing subjective clinical scores. In particular, sensor-based recordings of movement patterns in different patient cohorts are beneficial to deeply understand intra- and interday variability of motor impairments of chronic diseases.

This understanding is of major importance to characterize subgroups of patients or develop personalized approaches in order to evaluate disease progression and therapy response. The real-life data could also help detect changes early in the disease course, which is highly relevant for applying disease-stage-specific therapies. Future studies should evaluate the accuracy, usability as well as the technical and clinical validity of a sparse sensor setup in patients with movement disorders. This would set the basis for robust and valid sensor-based data recordings in real-life with the long-term goal to improve daily care and quality of life of patients using digital technologies.

In conclusion, our work shows that we can accurately perform a comprehensive sagittal-plane motion analysis with sparse sensor setups. We conclude that a comprehensive analysis including spatiotemporal, kinematic, and kinetic variables can best be performed with a sparse sensor setup that include sensors on the feet, the pelvis, and the thighs. We found that different setups performed better for different types of variables. A setup with feet and pelvis sensors was as accurate as the full setup for spatiotemporal and kinetic variables, while a setup with feet and thigh sensors was as accurate as the full setup for kinematic and kinetic variables, with a performance drop for the ankle moment up to 2 BW BH % using both sensor setups during running.

Therefore, when a comprehensive analysis is not necessary, the ideal sparse sensor setup is different for different applications. This study serves as a first step towards validating sparse inertial sensor sets for monitoring movements “in the wild”. Future validation studies should ensure that the processing between the IMU-based motion analysis and the ground truth motion analysis is as similar as possible, to avoid including processing differences in the results. This way, validity and usability of sparse sensor setups can be further evaluated for specific applications in order to allow the use of gait analysis in a wide spectrum of clinical and sports applications.

## Supporting information

S01: Averaged sagittal plane inertial sensor signals

S02: Individual simulation reports

## Footnotes

1 We use the notation [minimum value; maximum value] to show the range of the evaluation criteria.

