## Supplementary material for "Comparing sparse inertial sensor setups for sagittal-plane walking and running reconstructions": S01: Averaged sagittal plane inertial sensor signals

### Supplementary File S01: Averaged sagittal plane inertial sensor signals

Figure 1 and Figure 2 show the inertial sensor signals of the simulations and the reference signals for walking and running, respectively. The signals were averaged over all participants and speeds for each of the tested sensor setups. The sensor setups are abbreviated using the first letter of the segments equipped with a sensor: F-feet, S-shanks, T-thighs, P-pelvis. The setup FSTP corresponds to a full lower-body sensor setup with seven inertial measurement units (IMUs).

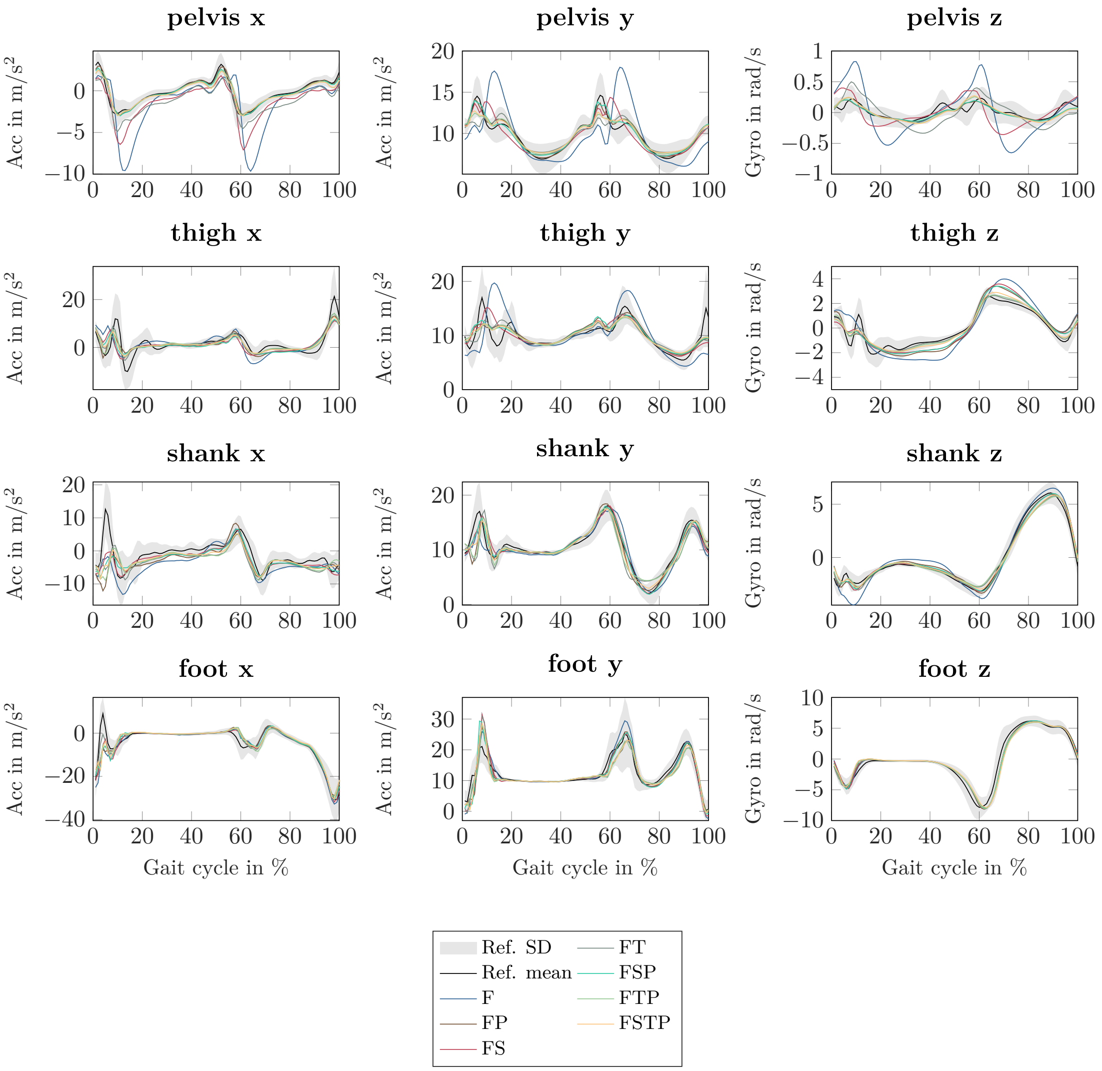


**Fig 1.** Sagittal-plane inertial sensor signals of the right lower limb for walking at all speeds, from the different inertial measurement unit (IMU) setups (colored lines) and the references values from optical motion capture system and force plate data (mean: black line, standard deviation: grey fill). All lines represent the mean over all participants from right heel strike to right heel strike.

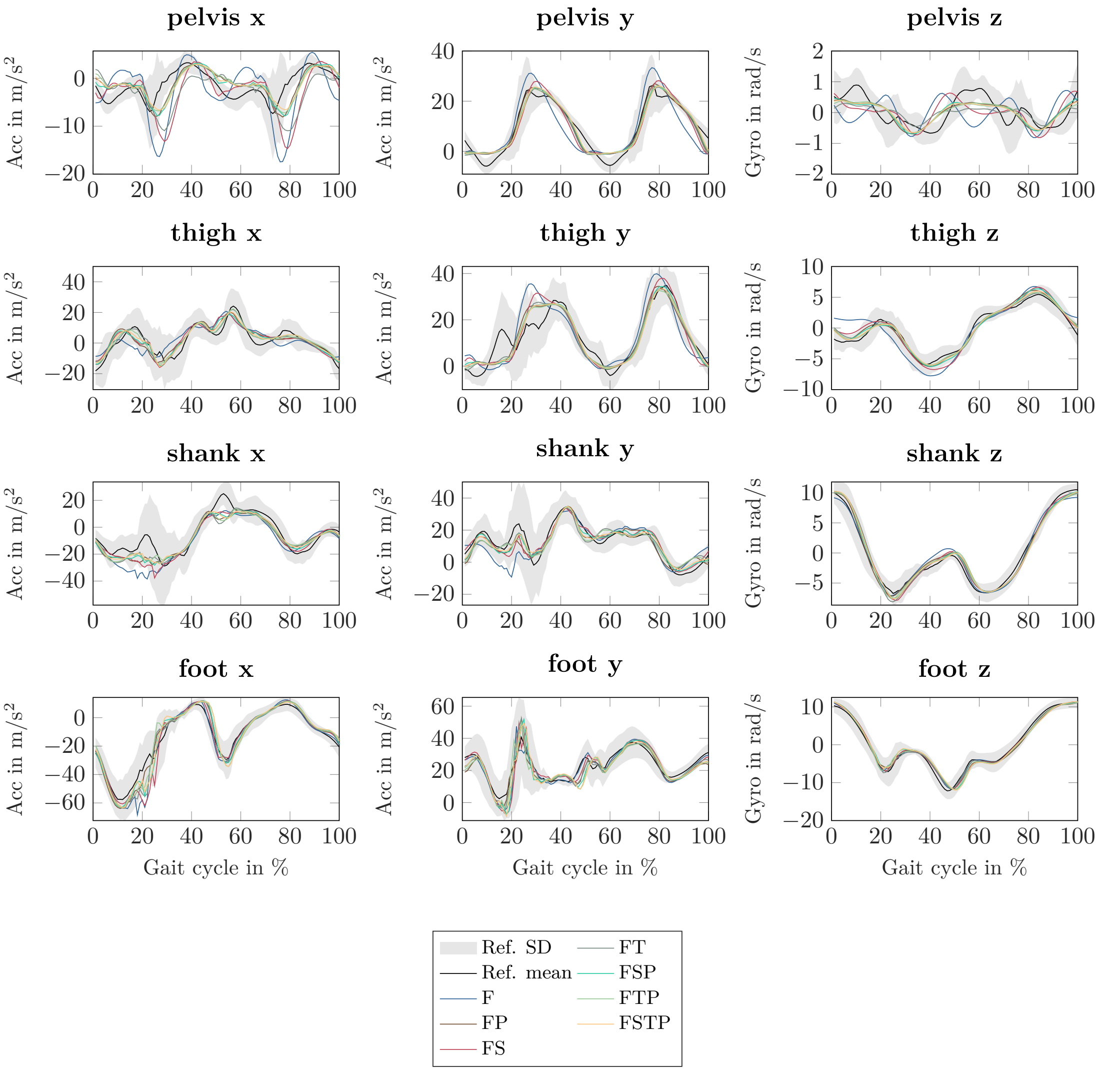


**Fig 2.** Sagittal-plane inertial sensor signals of the right lower limb for running at all speeds, from the different inertial measurement unit (IMU) setups (colored lines) and the references values from optical motion capture system and force plate data (mean: black line, standard deviation: grey fill). All lines represent the mean over all participants from left toe off to left toe off.
