## Supplementary material for "Comparing sparse inertial sensor setups for sagittal-plane walking and running reconstructions": S02: Individual simulation reports: P03_slowwalking_setup_FS_report.pdf

### Report of P03 slowwalking setup FS

June 10, 2024

#### 1 Solver

##### 1.1 Solver Status

- Status ID: Solve_Succeeded
- Status Message: Optimal Solution Found
- Number of iterations: 2931
- CPU time: 01:13:14 (HH:MM:SS)

##### 1.2 Solver Settings

- Solver: IPOPT
- tol: 0.0001
- max_iter: 20000
- constr_viol_tol: 0.001
- compl_inf_tol: 0.001
- acceptable_tol: 1e-06
- bound_frac: 0.001
- bound_push: 0.001
- hessian_approximation: limited-memory
- check_derivatives_for_naninf: no
- mu_strategy: adaptive
- linear_solver: mumps
- print_level: 5
- print_timing_statistics: yes
- For all other options, default values were used.

### 2 Problem

#### 2.1 General Information

- Model: Gait2dc
- Number of nodes: 100
- Symmetry: false
- Euler Method: BE
- Translation speed: 0.929 (m/s)
- Movement duration: 1.452 (s)
- Metabolic cost: 3.463 (J/m/kg)
- Objective Terms:

| name | weightedValue | weight | unweightedValue |
|---|---|---|---|
| regTerm | 9.921044e-04 | 1.000000e-05 | 9.921044e+01 |
| effortTermMuscles | 4.509230e+00 | 3.000000e+02 | 1.503077e-02 |
| trackAcc | 2.252410e+00 | 2.000000e+00 | 1.126205e+00 |
| trackGyro | 1.592152e+00 | 1.000000e+00 | 1.592152e+00 |

  tracked Variables:

  - Acc: foot_l, foot_r, tibia_l, tibia_r
  - Gyro: foot (left and right), tibia (left and right)

- Constraint Terms:

| name | normc |
|---|---|
| dynamicConstraints | 2.377781e-10 |
| periodicityConstraint | 3.845925e-16 |

#### 2.2 GRF

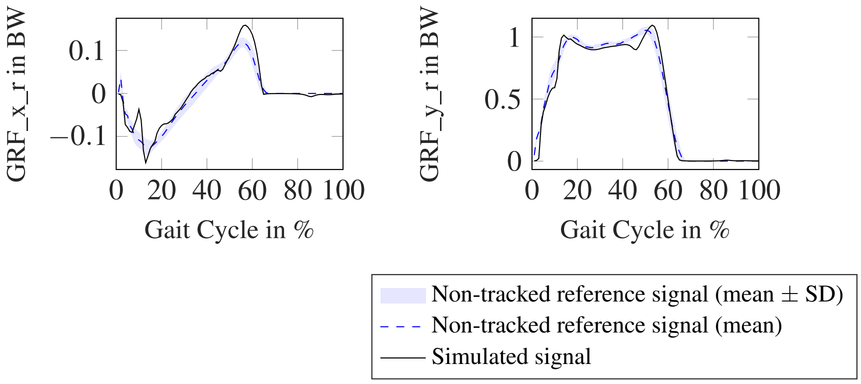

#### 2.3 acc

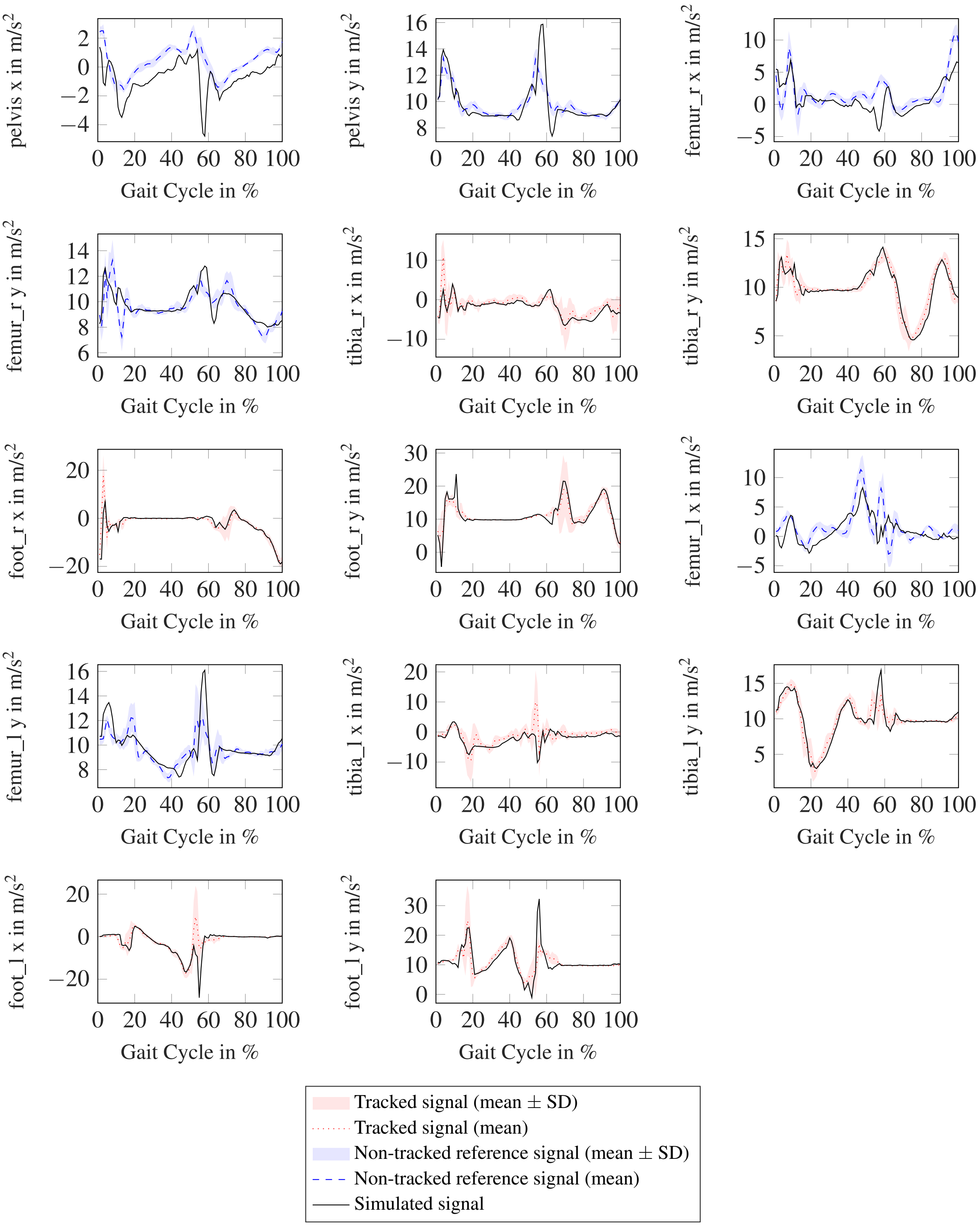
pelvis x in m/s$^2$
pelvis y in m/s$^2$
femur_r x in m/s$^2$
femur_r y in m/s$^2$
tibia_r x in m/s$^2$
tibia_r y in m/s$^2$
foot_r x in m/s$^2$
foot_r y in m/s$^2$
femur_l x in m/s$^2$
femur_l y in m/s$^2$
tibia_l x in m/s$^2$
tibia_l y in m/s$^2$
foot_l x in m/s$^2$
foot_l y in m/s$^2$
Gait Cycle in %
Tracked signal (mean ± SD)
Tracked signal (mean)
Non-tracked reference signal (mean ± SD)
Non-tracked reference signal (mean)
Simulated signal

#### 2.4 angle

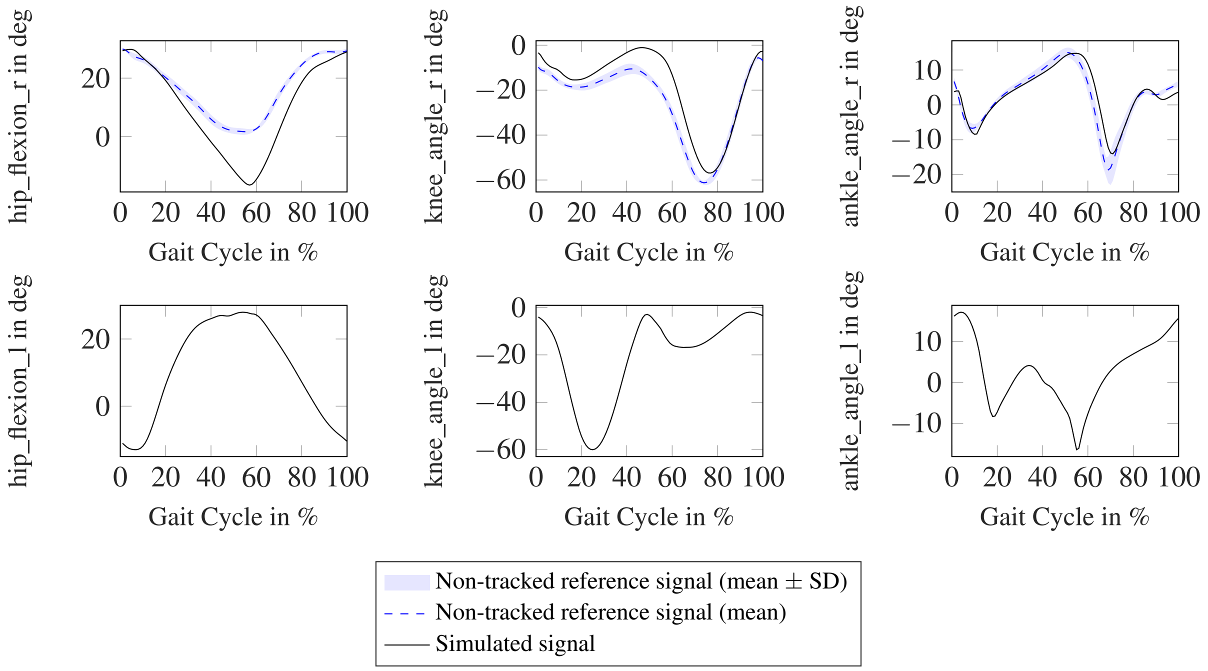


#### 2.5 gyro

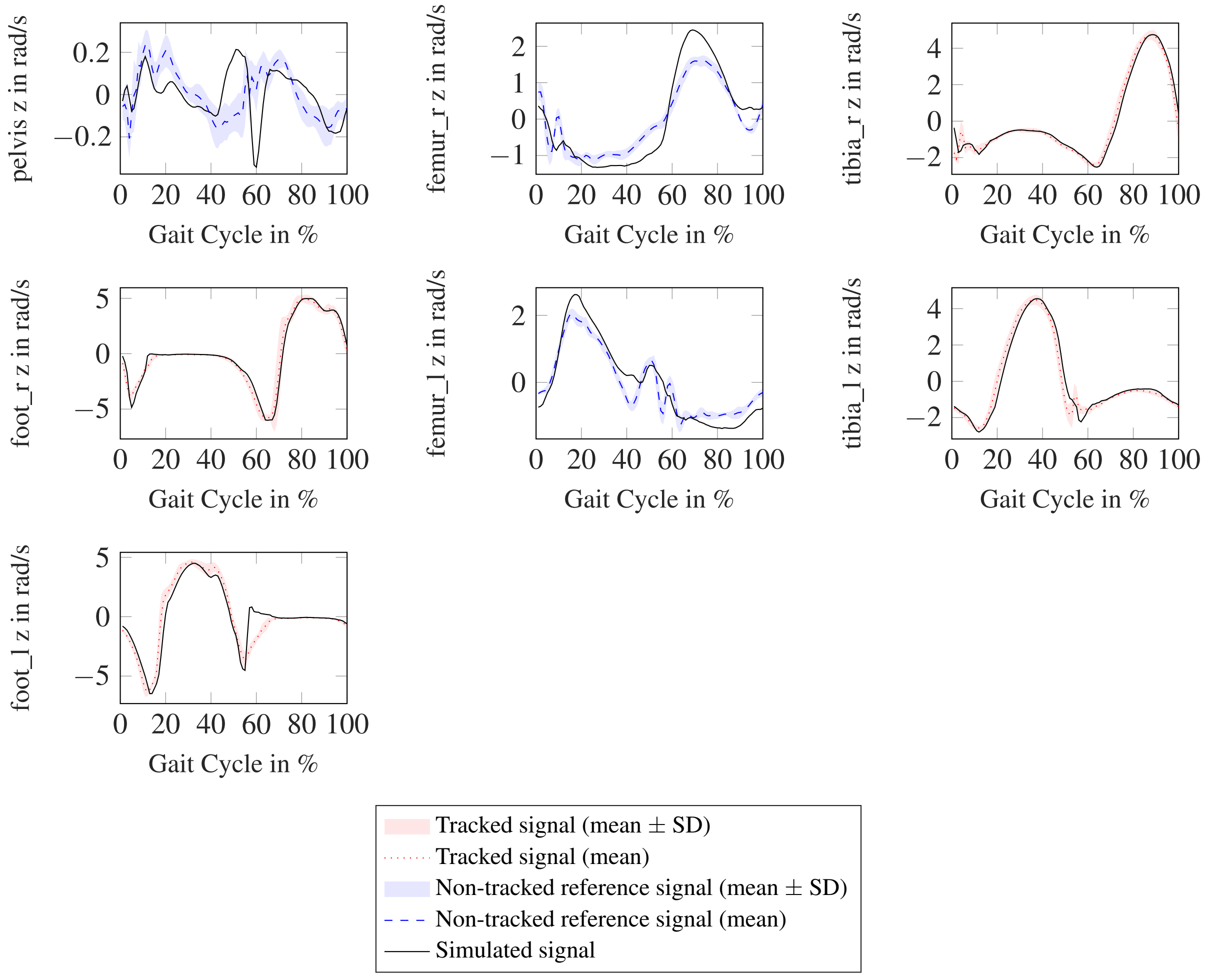

#### 2.6 moment

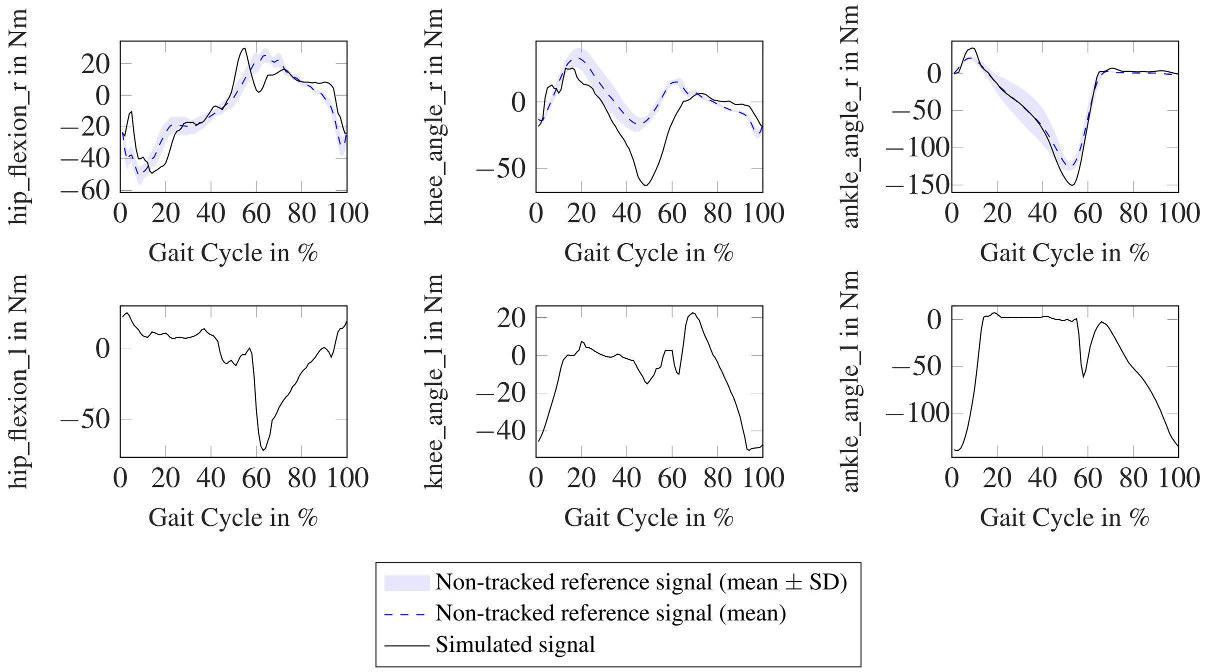


#### 2.7 muscleForce

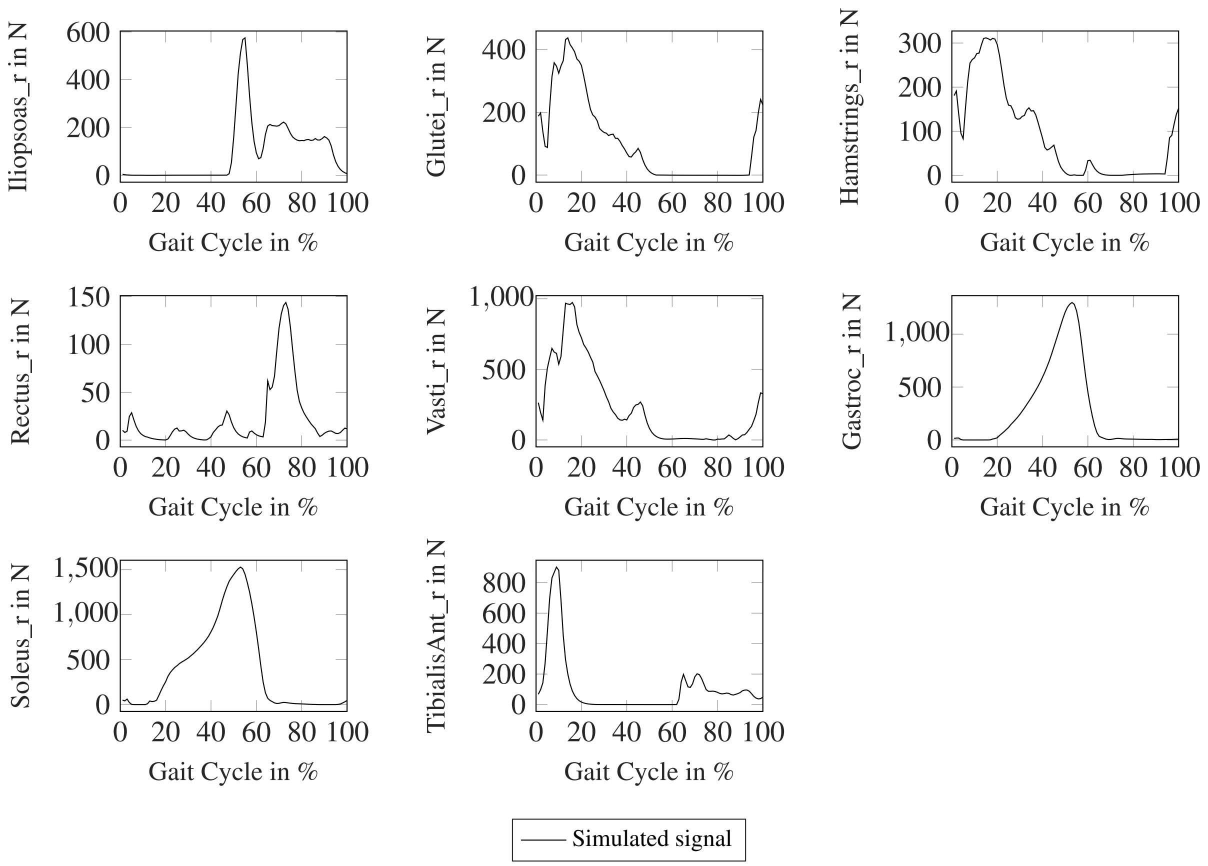
